# INTREPPPID - An Orthologue-Informed Quintuplet Network for Cross-Species Prediction of Protein-Protein Interaction

**DOI:** 10.1101/2024.02.13.580150

**Authors:** Joseph Szymborski, Amin Emad

## Abstract

An overwhelming majority of protein-protein interaction (PPI) studies are conducted in a select few model organisms largely due to constraints in time and cost of the associated “wet lab” experiments. *In silico* PPI inference methods are ideal tools to overcome these limitations, but often struggle with cross-species predictions. We present INTREPPPID, a method which incorporates orthology data using a new “quintuplet” neural network, which is constructed with five parallel encoders with shared parameters. INTREPPPID incorporates both a PPI classification task and an orthologous locality task. The latter learns embeddings of orthologues that have small Euclidean distances between them and large distances between embeddings of all other proteins. INTREPPPID outperforms all other leading PPI inference methods tested on both the intra-species and cross-species tasks using strict evaluation datasets. We show that INTREPPPID’s orthologous locality loss increases performance because of the biological relevance of the orthologue data, and not due to some other specious aspect of the architecture. Finally, we introduce PPI.bio and PPI Origami, a web server interface for INTREPPPID and a software tool for creating strict evaluation datasets, respectively. Together, these two initiatives aim to make both the use and development of PPI inference tools more accessible to the community.

**GRAPHICAL ABSTRACT:** 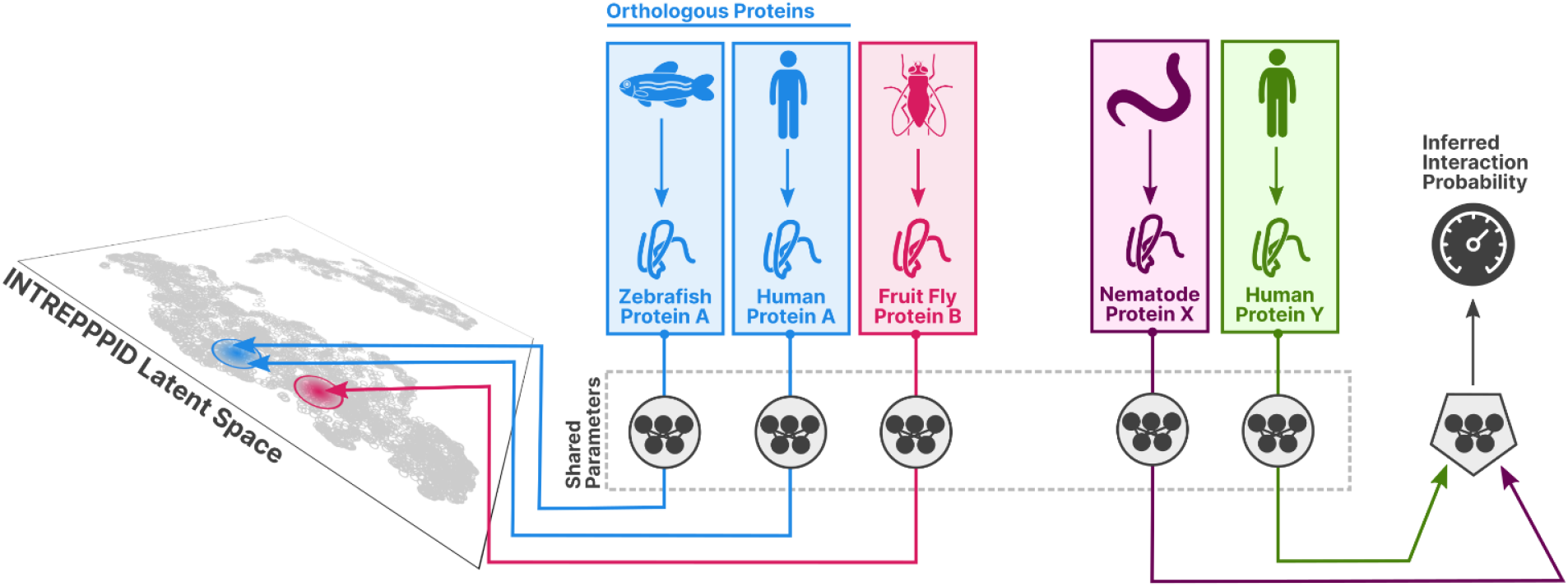

## INTRODUCTION

As the relevant technologies have improved, studies which experimentally verify protein-protein interactions (PPIs) have increased in size. In fact, multiple recent studies have published so-called “all-by-all” studies whereby the interaction of entire proteomes are experimentally evaluated. In September 2014, researchers assembled a reference network of approximately 14,000 interactions from a search space of 13,000 Human genes through a combination of yeast-two-hybrid (Y2H) assays and other validation experiments (1). Five years later, the same network (“HuRI”) expanded to approximately 53,000 interactions from a search space of 17,500 proteins (2).

While these studies provide an invaluable wealth of data, they represent a tremendous investment of time, labour, and money. A major consequence of the high cost of “all-by-all” studies is that it is prohibitively expensive to exhaustively perform them for all known species. In fact, it is currently only practical to limit these studies to a small number of well-studied model organisms. This is a problem common to many aspects of the life sciences, which we’ve taken to calling “the species gap”.

As of Frebruary, 2024 there are currently 793,217 taxa spanning 108,340 genera enumerated by the NCBI Taxon Database (3). Few, if any, experimental studies are capable of accounting for most, let alone all, of these taxa. For this reason, scientists have settled on a small pool of “model” organisms which exhibit desirable properties which lend themselves to a field of study and/or a certain methodology. Some properties, like relatively short life-cycles and small size, are favoured characteristics. These selection criteria further bias our already small sampling of all possible living subjects of study. To better demonstrate the extent of the species gap, we quantified the number of distinct organisms that are absent from the STRING Core PPI database (4), by calculating the intersection of taxa in this database and taxa in the NCBI Taxon Database. We found that only 30% of all taxa at the class rank and 16% of all taxa at the order rank were present in the STRING Core database (Figure 1), showcasing that we lack PPI information from the vast majority of species.

**Figure 1:**
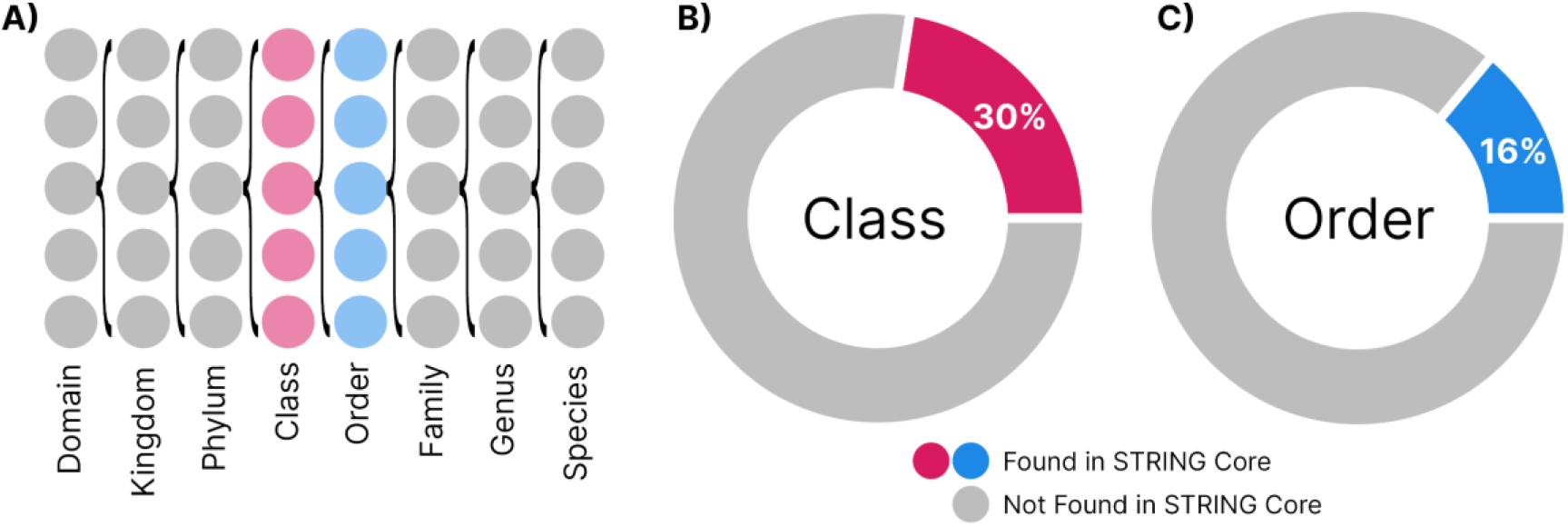
The species gap as illustrated by the STRING PPI database. (A) The hierarchy of biological taxa (5). (B-C) Doughnut charts indicating the proportion of all taxa represented in STRING Core at the class rank (B) and order rank (C).

*In silico* models of protein interaction have many favourable characteristics which may help narrow the species gap. Relative to Y2H or co-immunopercipitation assays, for example, which may take on the order of days and months to complete and analyze, *in silico* methods can take as little as seconds or minutes to complete. Furthermore, nearly all *in silico* PPI prediction methods operate entirely unattended, resulting in much lower labour costs. Finally, while some methods, like those which utilize deep learning, require initial hardware investments for training, inferring PPI prediction is often computationally inexpensive and can leverage commodity hardware. In summary, *in silico* methods and their low overall cost lend themselves naturally to the task of exhaustively searching protein-interaction spaces.

Practically, however, *in silico* methods commonly suffer from limitations that inhibit their use on less-commonly studied organisms. Namely, most *in silico* methods, especially those dependent on machine learning algorithms, require a training dataset comprised of protein interactions. These datasets must be sufficiently large as to provide a representative sample of interactions. They must also scale with the number of parameters being learned lest the model overfit. Other statistical methods which do not rely on machine learning also have similar requirements.

Regrettably, a major barrier to leveraging *in silico* methods for bridging the species gap outlined here is that lesser-studied organisms do not possess sufficient protein interaction data for which to train PPI inference models. In order for computational models to be able to help address the species gap, they must be able to make accurate PPI inferences on less-studied organisms, while using existing PPI data from well-studied organisms during training. Such “cross-species” inference requires that methods perform out-of-distribution predictions. Historically, PPI inference methods have been shown to fail to make accurate predictions on interactions that contain out-of-distribution proteins (6–8).

In 2012, a letter published by Park and Marcotte outlined the severity of a pervasive source of data leakage that was obscuring the inability of methods of the time to generalize to proteins present in inferred interactions that were out-of-distributions (6). The cause of the data leakage Park and Marcotte observed was the common “random splitting” approach to prepare cross-validation folds. Random splitting of interactions (between pairs of proteins) into folds allows for the same proteins appearing in both the training and testing sets, resulting in data leakage. Park and Marcotte found that when methods of the time were trained and tested on “strict” datasets where proteins present in the training set never appear in any of the other folds, most methods saw their testing accuracy drop to random levels. This precipitous drop in performance indicates that all the models tested had problems dealing with out-of-distribution proteins.

While Park and Marcotte found that out-of-distribution performance was poor across the landscape of methods available in 2012, recent studies have found that poor out-of-distribution performance still plagues current leading PPI inference methods. Studies by Dunham and Ganapathiraju in 2021 and Bernett *et al*. in 2023 corroborate Park and Marcotte’s 2012 findings (7, 8). In both studies, contemporary sequence-based PPI inference methods were tested on datasets that satisfy Park and Marcotte’s strictest validation schemes. Dramatic degradations in predictive power were observed in all cases, just as they were observed by Park and Marcotte in 2012. While *in silico* models for PPI inference have changed drastically since 2012 with the notable introduction and pervasiveness of deep learning methods, reported performance metrics of leading methods continue to be inflated as a result of the random-splitting of interactions (as opposed to stricter splitting based on proteins).

Furthermore, work by Hamp and Rost as well as Bernett *et al*. underscore an additional source of data leakage: distinct proteins with sequences which are nearly identical (8, 9). Amino acid sequences can be overwhelmingly redundant between distinct proteins. An important consequence of this redundancy is that even though no single protein appears in both training and testing sets, two proteins with all-but identical sequences may appear in both sets, thereby leaking data in the same way randomly-splitting PPI pairs does. The challenges presented by sequence similarity can be overcome by ensuring that proteins in the training set do not have a sequence identity above some set threshold with proteins in the testing set.

Some of the earliest methods devised to make inferences on protein interactions through sequence alone is Marcotte *et al*.’s *Rosetta Stone* method (10). The author’s outline a theory whereby interacting proteins of one species sometimes appear as fused homologous proteins in other species. Using this technique Marcotte *et al*. inferred interactions between proteins from *Escherichia coli* using information of proteins of other genomes which had similar sequences to the putative pair. The concept of “interologue” was soon after introduced to describe proteins whose interactions are conserved among their respective orthologues (11). Interologues allow us to extend some of our knowledge of interactions of one species to other species by identifying both sequences and interactions which are conserved between them (12, 13). These homology-based methods, while effective at identifying conserved interactions, have limitations related to unknown interactions, as acknowledged in the work that first introduced the term interologue: “Although potentially powerful, applying the interologue concept alone in estimating the significance of potential interactions precludes the finding of novel connections previously unidentified in other model organisms” (11).

Some of the first machine-learning methods for predicting protein-protein interactions were kernel-based methods operating on sequence features. The support vector machine (SVM) is an algorithm for finding the maximum-margin hyperplane for a classification or regression problem, relying on kernel transformations of the input data to solve non-linear problems (14). Effectively applying SVMs to the problem of PPI classification depends greatly on the design of the kernels used to represent protein pairs. These kernels are designed by researchers using expert-knowledge. These kernels may compare individual proteins to each other (“genomic kernels”) or pairs of proteins to other pairs (“pairwise kernels”). Furthermore, these kernels may compare a variety of features of the gene products being studied, including motifs, gene ontology (GO) annotations, sequence similarity, or some combination of these kernels (15).

As advances in hardware and related algorithms made research into Deep Neural Networks (DNNs) more feasible, they’ve proven to be effective in a number of problem domains, not least of which being PPI prediction. Richoux *et al*. demonstrated that twin neural-networks are far more efficient at PPI inference than other network architectures (16). Twin neural networks are models which consider pairs of inputs using the same trainable parameters rather than specifying parameters for each input, and are commonly used to make comparisons between two inputs (17, 18). This can result in networks with up to 50% fewer parameters, reducing the risk of overfitting and aiding few-shot performance (19). Most twin PPI prediction methods rely on convolutional neural networks (CNN) or recurrent neural networks (RNN) to encode feature sequences. The PIPR method by Chen *et al*., however, eschews these conventional architectures for a hybrid: the Recurrent Convolutional Neural Network (RCNN) (20). The RCNN architecture used in PIPR comprises of an initial CNN layer, whose output is pooled and then used as the input into a bidirectional Gated Recurrent Unit (GRU) RNN (21).

We’ve previously described RAPPPID, a method for PPI prediction that outperforms leading methods on strict validations sets that conform to the guidelines set out by Park and Marcotte (22). RAPPPID is a twin neural network which learns latent representations of sequences and inputs them into a classifier network to infer whether two proteins interact. RAPPPID achieves its high performance largely due to its adoption of AWD-LSTM and the Unigram tokenization algorithm as implemented by the SentencePiece software (23, 24). The former is a combination of regularization techniques that are appropriate and effective for long short-term memory (LSTM) RNNs. The input is tokenized using subword regularization, which helps regularize the network by sampling from a distribution of different segmentations of the input (25). Through the adoption of AWD-LSTM, Unigram tokenization, and balancing the number of parameters of the overall neural network, we were able to increase the generalization of the weights learned by RAPPPID.

Here, we introduce an extension of RAPPPID named INTREPPPID (**In**corporating **Tr**iplet **E**rror for **P**redicting **PPI**s using **D**eep Learning). INTREPPPID is specifically designed to improve cross-species performance with the aim of helping towards reducing the species gap. To achieve this, we designed INTREPPPID to learn latent representations of amino-acid sequences that are useful for predicting PPIs as well as identifying orthologues. Alongside RAPPPID’s original twin network for inference, INTREPPPID adds a triplet network which aims to minimize the Euclidean distance between latent representations of orthologous sequences while maximizing it for non-orthologous sequences. Parameters are shared between both the twin network and the triplet network, resulting in five networks with a single set of shared parameters, which we call a “quintuplet” network. Because of INTREPPPID’s architecture, its latent space encodes both information relevant to PPI inference as well as orthology. This allows us to enforce an inductive bias loosely based on the concept of interologues.

Our analysis shows that INTREPPPID outperforms all leading methods tested on both intra-species and cross-species PPI predictions. Further, we demonstrate that INTREPPPID effectively learns the orthology of proteins. Finally, we make public to the community PPI.bio and PPI Origami. PPI.bio is an open-source interface for INTREPPPID and RAPPPID hosted on the internet. This allows researchers to infer PPIs without installing any specialized software or using the command-line. PPI Origami is a software package for creating reproducible strict PPI databases in an automated fashion. By automating the dataset generation process, we hope to reduce common sources of data leakage in PPI datasets used for PPI inference.

## MATERIALS AND METHODS

### The INTREPPPID architecture

INTREPPPID integrates RAPPPID’s twin neural network for the PPI Classification task with a triplet neural network used for a novel Orthologous Locality task. Learning the two networks jointly results in five encoder layers with shared parameters in what we’ve come to call a quintuplet network architecture.

INTREPPPID uses the same AWD-LSTM encoder as the one used in RAPPPID to learn vector representations of protein sequences. This encoder is a regularized recurrent neural network which has been shown to improve perfomance on tasks that require inferring interactions between proteins absent from the training set (22). Before encoding, sequences are first tokenized using the Unigram algorithm as implemented by the SentencePiece library (24, 25). INTREPPPID uses the vector representations generated by the encoder for two downstream tasks: the *orthologous locality* task and the *PPI classification* task. The encoder as well as the two tasks are trained in an end-to-end fashion.

INTREPPPID’s quintuplet architecture means that training the network relies on additional orthology data. At inference time, however, only the protein classification twin network is used, so no information other than the amino acid sequences of the proteins in question are required as input.

#### The PPI classification task

INTREPPPID infers PPIs by learning latent representations of the candidate protein sequences which can be classified by a fully-connected neural network outputting the interaction probability. The two amino acid sequences (**s**_**β**_ and **s**_**γ**_) are first tokenized into vectors **x**_**β**_ and **x**_**γ**_ using the Unigram algorithm. The Unigram algorithm segments the amino acid of the protein sequence into groups using a vocabulary of 250 tokens. The Unigram algorithm also employs subword regularization. Unigram draws tokens for a given sequence from a pobabilistic distribution of all possible combination of tokens in the vocabulary, meaning that the same sequence can be tokenized differently between successive calls to the tokenizer.

Once the tokenized vectors **x**_**β**_ and **x**_**γ**_ are obtained, they are inputted into an encoder (*e*(⋅)) which outputs a latent vector for each protein (**z**_**β**_ and **z**_**γ**_, respectively):

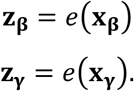

The encoder has an Averaged Weight-Decayed Long Short-Term Memory (AWD-LSTM) architecture, that leverages drop-connect and averaged optimization (here, stochastic weight averaging), among other things, to regularize and improve generalization (see RAPPPID’s study for details (22)). The parameters of the encoder that infers **z**_**β**_from **x**_**β**_are the same as those of the encoder that infers **z**_**γ**_ from **x**_**γ**_, making the PPI classification network a twin neural network.

Finally, the latent vectors **z**_**β**_ and **z**_**γ**_ are averaged and inputted into a classifier network *g*(⋅) which outputs an interaction probability ŷ:

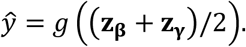

The classifier *g*(⋅) is a two-layer, fully-connected neural network whose hidden layer is activated by the Mish function (26). The fully-connected neural network outputs a single real number which, once activated by the sigmoid function, represents the probability that the proteins represented by the amino acid sequences **s**_**β**_and **s**_**γ**_ interact. This interaction probability as well as the true label are inputted into the binary cross-entropy loss to obtain a loss for the PPI classifcation network:

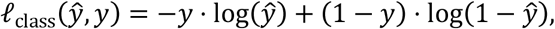

where *y* is a binary label representing whether the two proteins truly interact (*y* = 1) or not (*y* = 0).

#### The Orthologous Locality task

INTREPPPID introduces a novel task to help inform the PPI classification network. The orthologous locality task is to learn a latent space whereby the Euclidean distance between latent representations of proteins which are orthologous to one another are within some margin *m*. Conversely, non-orthologous proteins are to have a Euclidean distance greater than *m*.

To achieve this, we adopt a triplet network and a triplet margin loss *l*_ortho_ following the description of (27):

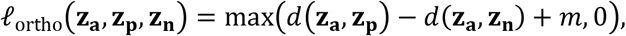

where **z**_**a**_ is the latent vector of some “anchor” protein for which there is a known orthologous protein with a latent vector **z**_**p**_, **z**_**n**_ is the latent vector of a randomly chosen protein which is not an orthologue of **z**_**a**_, *m* is a margin factor and *d* is the Euclidean distance:

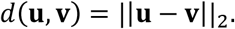

For the experiments conducted herein, the margin *m* was fixed at a value of 1.

The principle of the triplet network is to, at every training step, maximize the distance between two non-orthologous proteins (the “anchor” and “negative” proteins), as well as minimize the distance between two orthologous proteins (the “anchor” and “positive” proteins).

The anchor protein, with amino acid sequence **s**_**a**_, can be any protein in the dataset for which orthology data is known. All orthology data for the experiments conducted herein are from the OMA database of orthologues (28). In the event that orthologues are known for the protein which **s**_**β**_ encodes, **s**_**β**_ is chosen to be **s**_**a**_.

The positive and negative proteins (with amino acid sequences **s**_**p**_ and **s**_**n**_, repectively) are chosen relative to **s**_**a**_. **s**_**p**_ is chosen to be the sequence of a randomly chosen protein who, according to the OMA database, belongs to the set of sequences of proteins orthologous to the anchor protein. At every training step, **s**_**p**_ is drawn randomly from this set of proteins orthologous to **s**_**a**_, meaning that the positive protein for a given anchor protein may not be the same between training steps or epochs. The negative protein is sampled in a similar manner to the positive protein, but rather than being sampled from the set of proteins orthologous to **s**_**a**_, it’s sampled from the set of proteins not orthologous to **s**_**a**_. The amino acid sequence of the anchor, positive, and negative proteins (**s**_**a**_, **s**_**p**_ and **s**_**n**_) are tokenized and encoded in the same manner as **s**_**β**_ and **s**_**γ**_, resulting in their latent representations **z**_**a**_, **z**_**p**_, and **z**_**n**_.

Optimizing the triplet margin loss in this manner maximizes the Euclidean distance between latent vectors of proteins that are not orthologues, while minimizing the distance between vectors of orthologues. Our adoption of the triplet margin loss follows evidence that such a loss is an effective way of creating feature descriptors that are “spread-out” over the latent space, which in turn increases performance on downstream classification tasks (27, 29). Furthermore, orthologues typically share the same protein functions and therefore offer a good inter-species basis for the downstream PPI classification task (30, 31).

### INTREPPPID’s loss function

The final INTREPPPID loss function that is optimized at training time is the sum of *l*_ortho_ and *l*_class_,

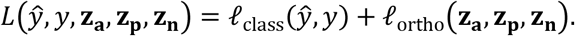

In this way, INTREPPPID optimizes both the orthologous locality and PPI classification losses simultaneously, in an end-to-end fashion, ensuring that latent vectors simultaneously encode orthology and PPI information.

### Preparing strict PPI datasets

Experiments were run using datasets with strict data splitting folds, described as “C3” by Park & Marcotte (6). Namely, we ensured that there was no overlap of proteins between any of the training, validation, or testing datasets. Additionally, we controlled for sequence similarity of proteins between the training, validation, and testing sets which is a known vector for information leakage (8, 9). Supplementary Table S1 outlines summary statistics for the datasets used in this study. The formation of these strict datasets is described below.

A PPI dataset 𝒟 is a set of interacting protein-pairs reflecting binary interactions *D*_*i*_ ∈ 𝒟. 𝒟 is partitioned into three mutually disjoint subsets: a training subset (𝒟_Tr_), a testing subset (𝒟_Te_), and a validation subset (𝒟_V_). Let *P*_Tr_, *P*_Te_, and *P*_V_ be the set of proteins present in 𝒟_Tr_, 𝒟_Te_, and 𝒟_V_, respectively, and let 𝒫 be the collection of protein sets {*P*_Tr_, *P*_Te_, *P*_V_}.

We define a strict PPI dataset as a dataset 𝒟 (with its collection of protein sets 𝒫) that fulfills the following two criteria.

#### Criterion 1 - Distinct Protein Identity

The resulting protein sets *P*_Tr_, *P*_Te_, *P*_V_ of the dataset must be mutually disjoint:

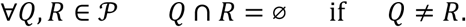

#### Criterion 2 - Distinct Sequence Identity

The amino acid sequence of any pair of proteins belonging to different protein subsets must have a sequence identity no greater than 90%:

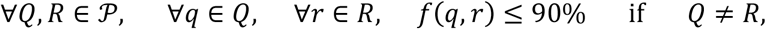

where *f*(⋅,⋅) is some protein sequence identity function.

For the datasets used in the experiments carried out herein, the UniRef90 identifiers provided by UniProt were used to determine sequence identity, and as such we adopt the same definition of sequence identity (32, 33).

#### High-confidence examples of PPIs

To minimize our exposure to falsely labelled PPIs, we required a database that provided information regarding the confidence of the interactions. We selected the STRING database as the source of PPIs for our datasets as it is large, contains information for many species, and integrates confidence scores from a number of modalities (4). We exclude all PPIs which do not have a combined confidence score greater or equal to 90%, and only include PPIs which are classified as “physical interactions”. PPI information for seven species were included in our analyses (Supplementary Table S2 in File S1).

#### Negative examples of PPIs

As INTREPPPID is a supervised method, it requires examples of pairs of proteins which do *not* interact (negative examples), in addition to pairs which do (positive examples). Obtaining negative examples of PPIs is a fraught process as there exists no one agreed upon “gold standard” of proteins which do not interact with one another. While many approaches for obtaining negative PPI examples have been devised over the years, evidence has shown that many of these methods introduce a great amount of bias (34). For this reason, we opted to sample pairs of proteins in a uniformly random manner to represent our negative examples. For the training set, negatives were randomly drawn without replacement from the set *N*_Tr_, defined as:

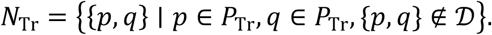

Negative examples for the testing and validation sets were drawn from *N*_Te_ and *N*_V_, which were similarly defined as the set of protein-pairs of *P*_Te_ and *P*_V_, respectively, which were not in 𝒟 (*i*.*e*., were not considered interacting in the dataset).

Random sampling of protein-pairs requires the fewest assumptions about the distribution of negative pairs, thereby reducing bias, albeit at the risk of false-negatives. Through the exclusion of known interactions from *N*_Tr_, however, we’ve reduced the likelihood of drawing false-negatives.

#### Considerations for cross-species datasets

For experiments herein, it was necessary to prepare datasets suitable for testing the cross-species task. Namely, we needed to construct training sets whose PPIs were composed of proteins of one species, and many corresponding testing sets each belonging to a different species. All combinations of the training set and the many testing sets must satisfy both Criteria 1 and 2. The first criterion is typically met *de facto*, as protein identifiers between species normally do not overlap. In order for the second criterion to be met, positive and negative interactions of proteins that do not satisfy Criterion 2 were excluded from the datasets.

### Generating strict PPI datasets with PPI Origami

We have written a command-line interface (CLI) tool named “PPI Origami” with the aim of making the task of preparing strict PPI datasets in a replicable and automated fashion. PPI Origami has three primary functions: *downloading* primary databases, *processing* primary databases into strict datasets, and an *analysis* function.

PPI Origami allows for the downloading of primary databases as an integrated part of its pipeline. By directly handling downloading third-party databases, PPI Origami ensures that different users of the software are downloading the same files in the same way. PPI Origami also stores downloaded files in a distinct location from the processed files, isolating primary files from accidental changes.

PPI Origami’s main function is to process and transform primary databases into strict PPI datasets. It can create datasets that satisfy both criteria of strict PPI datasets, filtering low-confidence positive examples, and drawing negative examples as previously described.

PPI Origami allows for simple reporting of basic statistics of PPI datasets it generates. These reports allow for convenient verification of whether datasets meet both criteria of strict datasets, as well as other statistics of interest such as the ratio of interactions between dataset cross-validation subsets.

We hope that PPI Origami encourages other researchers to incorporate strict PPI datasets in the future development and analyses of supervised PPI inference methods.

### Training Procedure

INTREPPPID was trained using the Ranger21 optimizer, which uses AdamW with a look-ahead mechanism (44, 45). An “explore-and-exploit” learning rate scheduled was used, with a maximum learning rate of 0.01 (46). INTREPPPID was trained for 100 epochs. The weights of the epoch with the lowest validation loss were used in the analyses herein (and for evaluation on the test set). Hyperparameters used to train INTREPPPID were informed by those selected during the development of RAPPPID, and are listed in Supplementary Table S3.

## RESULTS

### INTREPPPID utilizes an orthology-informed quintuplet network for PPI prediction

INTREPPPID is a deep learning model for inferring the interaction of two proteins given their amino acid sequences and is designed to improve the performance of PPI inference of proteins from species other than the ones on which it is trained. It is an extension of our previous work RAPPPID, a twin neural network PPI inference method, which consistently outperformed other methods on strict evaluation datasets (22). INTREPPPID adopts a quintuplet architecture, which ensures its latent space encodes orthology data (Figure 2A, Methods). Its architecture consists of two components: a triplet network responsible for the *orthologous locality* task and a twin network for the *PPI classification* task. The encoders of these networks share weights and parameters, forming a quintuplet architecture.

**Figure 2:**
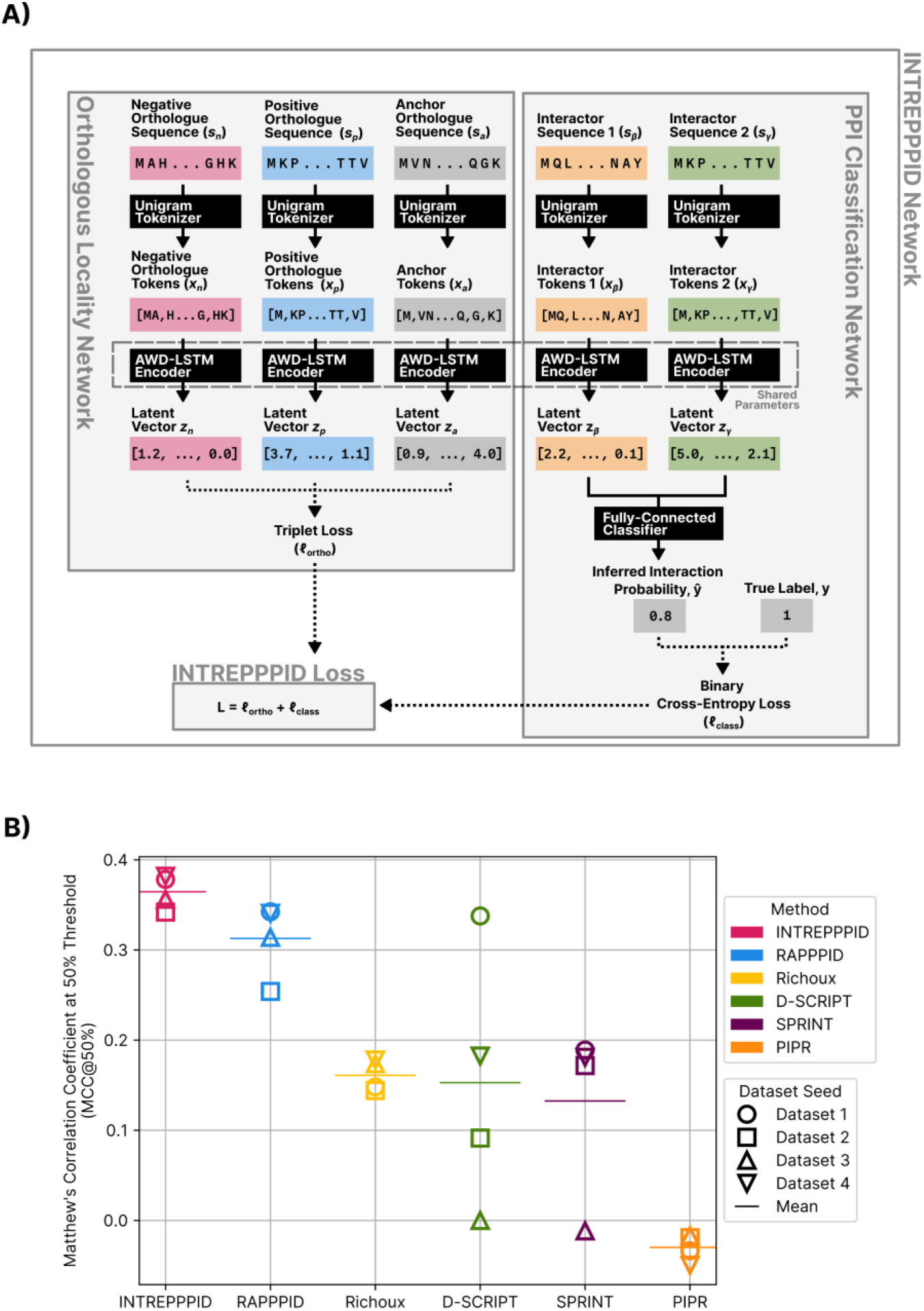
INTREPPPID’s architecture and performance. **(A)** A schematic overview of INTREPPPID’s quintuplet architecture. The leftmost panel depicts the triplet network responsible for the orthologous locality task, while the rightmost panel depicts the triplet network responsible for the PPI classification task. During training, INTREPPPID learns both tasks jointly through an overall loss function, which is the sum of the losses of the two tasks. **(B)** Test (C3) Matthew’s Correlation Coefficient (MCC) of leading PPI inference methods trained and tested on *H. sapiens* datasets.

INTREPPPID infers PPIs by learning latent representations of the candidate protein sequences. The two amino acid sequences are first tokenized into two vectors using the Unigram algorithm as implemented by the SentencePiece algorithm (24, 25). Then, they are provided as input to Averaged Weight-Decayed Long Short-Term Memory (AWD-LSTM) encoders, a type of regularized recurrent neural networks with the ability to make long-term connections. The obtained latent embeddings are then inputted into a classifier network which outputs an interaction probability. The orthologous locality task is introduced so that the latent representations of orthologous proteins are close to each other, while the non-orthologous proteins have distant representations (as measured using Euclidean distance). This auxiliary task helps regularize the PPI classification network by imposing biologically relevant constraints on the latent space. This is achieved using a triplet network with a triplet margin loss. As input, this network receives the tokenized vectors of two orthologous proteins (used as “anchor” and “positive” proteins) and a third “negative” protein that is not orthologous to the anchor protein. The tokenized vectors of these proteins are provided to AWD-LSTM encoders with identical architecture and weights to those of the twin network. In each training step, the model maximizes the distance between two non-orthologous proteins while minimizing the distance between two orthologous proteins. During training, both PPI Classification and Orthologous Locality networks are trained jointly in an end-to-end fashion. During inference time, only the twin PPI classification network is used.

In this study, we used orthology information from the OMA database (28) and the PPI interactions from the STRING database (4). These datasets were carefully processed to form strict C3 datasets (6) for training and evaluation within and across species. Moreover, we have developed a command-line interface (CLI) tool, “PPI Origami”, to automatically download primary PPI databases, and process them into strict PPI datasets. Methods describes the datasets, preprocessing steps, PPI Origami, and training procedure.

### INTREPPPID outperforms leading models for prediction of human PPI

First, we set out to evaluate INTREPPPID’s performance in an intra-species PPI prediction task: trained and tested on *H. sapiens* interactions. Table 1 shows the performance of INTREPPPID against five leading PPI prediction methods. These methods include RAPPPID, our previous method upon which INTREPPPID expands, D-SCRIPT, a method designed for the cross-species task, PIPR, a twin network with an RCNN encoder, SPRINT, a sequence-similarity-based method, and the twin networks proposed by Richoux *et al*. (16, 20, 22, 35, 36). INTREPPPID performed slightly better than RAPPPID across all metrics, and outperformed all other methods (Table 1, Figure 2B). Richoux *et al*., D-SCRIPT, and SPRINT perform similarly on average, with D-SCRIPT exhibiting a particularly high degree of variability across the different datasets tested. We observed severe over-fitting while training and testing PIPR on all datasets.

**Table 1:** Average performance of five leading PPI inference methods trained and tested on four H. sapiens PPI datasets. The performance metrics are calculated on strict C3 test sets (see Methods). A higher value for the first four metrics and a lower value for the last metric (Brier Score) shows a better performance. Bold-face indicates the best value. The following abbreviations are used. AUROC: Area under the receiver operating characteristic; AP: Average precision; MCC: Matthew’s Correlation Coefficient, SD: Standard deviation.

| Method | AUROC<br>Mean<br>(±SD) | AP<br>Mean<br>(±SD) | MCC@50%<br>Mean<br>(±SD) | F1@50%<br>Mean<br>(±SD) | Brier Score<br>Mean<br>(±SD) |
| --- | --- | --- | --- | --- | --- |
| INTREPPPID | **0.752**<br>(±0.017) | **0.730**<br>(±0.022) | **0.365**<br>(±0.067) | **0.697**<br>(±0.019) | **0.200**<br>(±0.004) |
| RAPPPID | 0.732<br>(±0.024) | 0.717<br>(±0.015) | 0.313<br>(±0.041) | 0.630<br>(±0.025) | 0.215<br>(±0.006) |
| D-SCRIPT | 0.634<br>(±0.125) | 0.627<br>(±0.086) | 0.153<br>(±0.144) | 0.506<br>(±0.282) | 0.402<br>(±0.116) |
| Richoux et<br>*al.* | 0.606<br>(±0.009) | 0.614<br>(±0.015) | 0.161<br>(±0.017) | 0.370<br>(±0.030) | 0.274<br>(±0.011) |
| SPRINT | 0.597<br>(±0.061) | 0.641<br>(±0.064) | 0.133<br>(±0.096) | 0.566<br>(±0.048) | 0.493<br>(±0.003) |
| PIPR | 0.482<br>(±0.006) | 0.484<br>(±0.003) | -0.030<br>(±0.015) | 0.318<br>(±0.010) | 0.487<br>(±0.005) |

### INTREPPPID maintains high cross-species performance on multiple organisms

INTREPPPID was designed to improve on RAPPPID’s ability to make predictions across species. We therefore sought to compare INTREPPPID’s cross-species performance with that of RAPPPID’s as well as other leading PPI prediction methods.

When trained on *H. sapiens* and tested on various species (C3), ranging from *Mus musculus* (house mouse) to *Arabidopsis thaliana* (thale cress), INTREPPPID predominantly outperformed other methods (Figure 3A, Supplementary Table S4), with only RAPPPID rarely surpassing its performance in some metrics. Figure 3A shows the mean AUROC (across 3 datasets) of different methods, with INTREPPPID significantly outperforming all other methods (one-sided Wilcoxon signed rank test vs. RAPPPID = 0.03). On the cross-species task, the performance of PIPR and SPRINT were close to a random classifier. Both D-SCRIPT and the Richoux *et al*. method performed similarly to one another, but consistently scored below RAPPPID and INTREPPPID.

**Figure 3:**
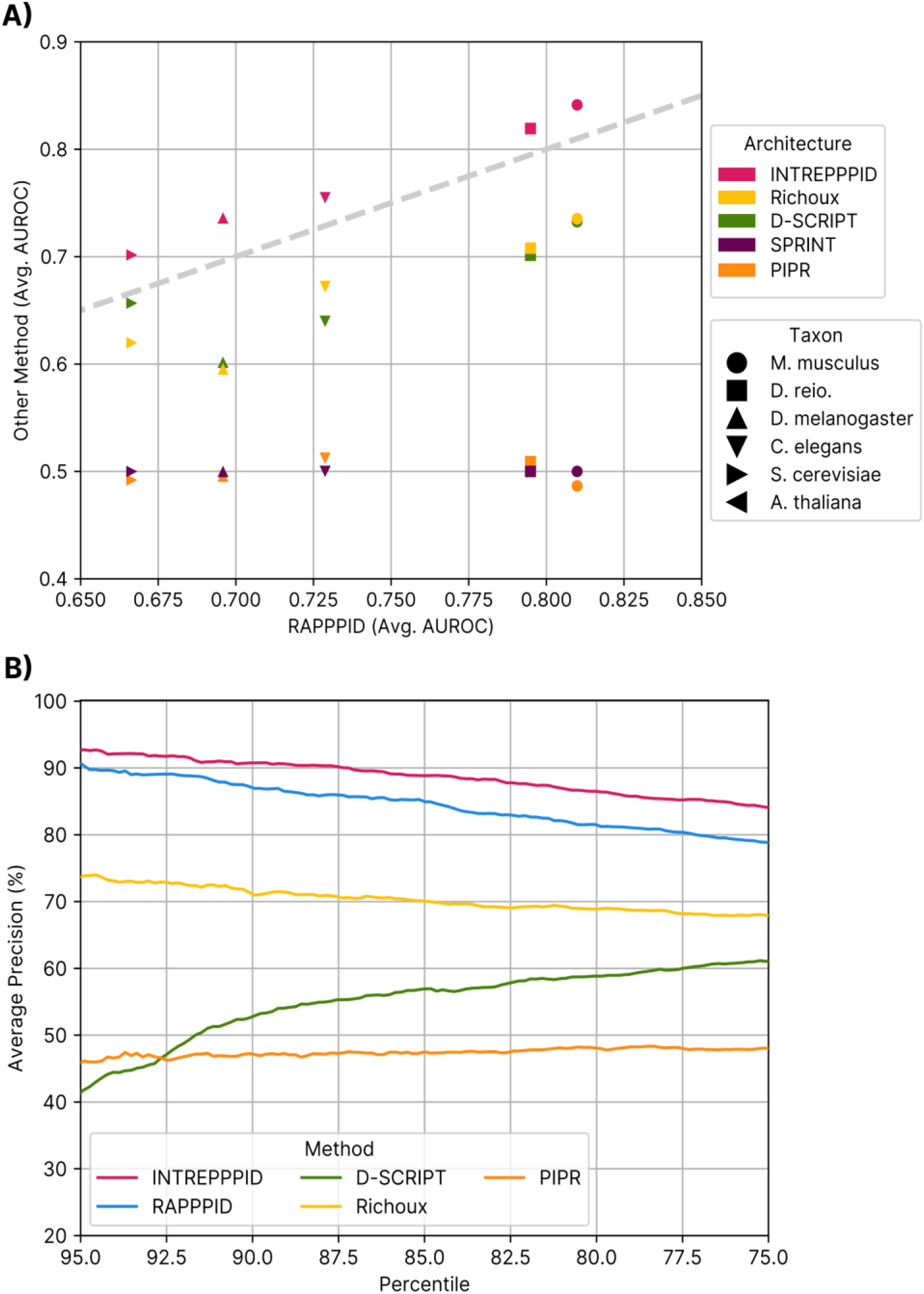
**(A)** Average cross-species AUROC of leading PPI inference methods. The y-axis shows the AUROC of each method (shown as different colours) against the AUROC of RAPPPID (x-axis). Each model was trained on H. sapiens and tested on various species (indicated by the different markers). Markers above the dashed grey line indicate instances where a method outperformed RAPPPID, while markers below the line indicate under-performance relative to RAPPPID. **(B)** The precision of inferred interactions among the *k*^th^ percentile of the scores inferred by the method indicated by the line colour. The x-axis shows *k* and the y-axis shows and precision along the y-axis. Interactions were inferred for datasets from seven different species. The mean across these datasets is reported by the solid line.

In many applications, one is particularly interested in the precision of a model among the highest-ranking predicted edges (*e*.*g*., to validate them experimentally). To evaluate INTREPPPID’s suitability for such tasks, we conducted an experiment whereby we measured the precision of predictions whose scores are above various percentiles (Figure 3B). Considering the average across all species tested, INTREPPPID achieved a precision greater than 90% among predictions above the 85^th^ percentile, and even when we considered the 75^th^ percentile, its mean precision remained above 85%. These observations confirm that INTREPPPID is suitable for applications where very high precision is paramount.

### INTREPPPID learns protein representations that exhibit strong orthologous locality

Through its orthologous locality task, INTREPPPID learns latent representations of orthologues which are near to one another in the Euclidean space, while ensuring latent representations of non-orthologous sequences remain distant. To qualitatively assess the locality of orthologues in latent space, we calculated the embeddings of the sequences of 1,000 randomly sampled proteins present in the OMA database of orthologues and projected them onto a two-dimensional space using the UMAP algorithm (Figure 4A)(37). The projections show that latent embeddings of orthologues cluster tightly in INTREPPPID’s latent space, while embeddings from RAPPPID (which has a similar architecture, but without the triplet network) do not.

**Figure 4:**
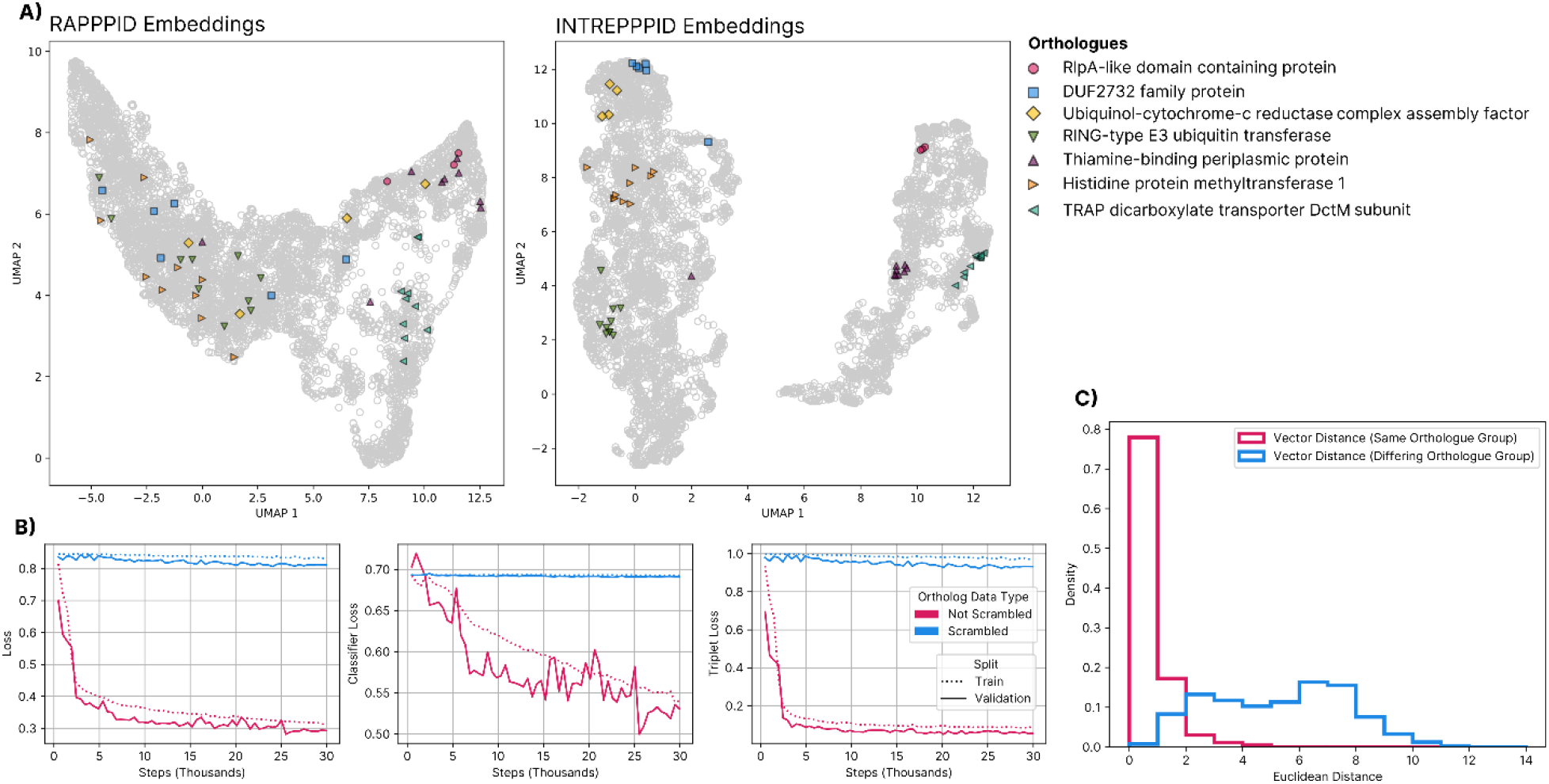
The effect of orthologous locality task. **(A)** A random sample of protein sequences from various species were embedded using RAPPPID (left) and INTREPPPID (right) and projected onto a 2D space using the UMAP algorithm. Distinct markers indicate proteins which belong to the same orthologous group according to the OMA database. The orthologous groups shown were chosen at random. **(B)** Training (dotted line) and validation loss (solid line) of INTREPPPID trained on *H. sapiens* datasets whose orthologue data is “scrambled” (blue) and “not scrambled” (magenta). The overall loss (left), the loss of the classification network (middle), and the loss of the triplet network (right) are plotted above, demonstrating that the “scrambled” dataset fails to optimize any aspect of the loss in any appreciable way. **(C)** Histogram of the Euclidean distance between INTREPPPID latent vector representations of protein sequences of the same orthologous group (magenta) and of differing orthologous groups (blue).

A more quantitative analysis of embedding distances further confirmed that INTREPPPID’s latent space exhibits strong orthologous locality. Namely, the Euclidean distance between latent representations of orthologues was on average 0.822, while the distance between the latent representations of non-orthologous proteins is on average 5.31 (Figure 4C). Nearly 80% of INTREPPID’s latent representations of orthologues fell under the triplet loss margin of 1, while virtually all non-orthologues had a distance greater than the margin between them.

Together, these analyses confirm that INTREPPPID successfully accomplishes the orthologous locality task.

### INTREPPPID’s performance gain is due to the biologically-relevant information encoded by orthology

To further assess the role of orthology information on the performance of INTREPPPID, we conducted an experiment in which we trained INTREPPPID on two separate datasets: one constructed using orthologue data from the OMA database (called “not scrambled”), and one where *random* pairs of proteins from the OMA database were labelled orthologues (“scrambled”). After training and testing on both variants of the H. sapiens dataset, we found that INTREPPPID’s performance dropped to near random in the “scrambled” case (Supplementary Table S5, AUROC = 0.524). Moreover, on this dataset, neither the classification loss nor the triplet loss significantly changed over 30,000 training steps (Figure 4B).

INTREPPPID’s low testing performance on “scrambled” data suggests that the mere exposure of sequences from various species is not sufficient to increase PPI classification performance. The impact of the “scramble” dataset on both the classification and triplet losses further suggests that the orthologous locality task is complimentary to the PPI classification task. Ablating the biological meaning of the orthology data conversely results in the orthologous locality task having a deleterious effect on the PPI classification task. These results show that the encoding of orthology information increases INTREPPPID’s performance due to its biological relevance.

### A web server and interface for RAPPPID & INTREPPPID

While we have made best efforts to make RAPPPID and INTREPPPID as simple as possible to run on a local computer with minimal computational requirements, it currently requires knowledge of the command-line interface, the Python package installer (pip), and for some use-cases, the Python computer language. These prerequisites, while often quite accessible to researchers with relevant programming experience, can be too great a hurdle for others for whom RAPPPID and INTREPPPID might otherwise be of great use.

To address this, we have created a website which acts as a graphical interface for a server which is capable of performing proteome-scale inference in about fifteen minutes (https://ppi.bio). The source code that runs PPI.bio is available publicly on GitHub and can be found by visiting the URI: https://github.com/Emad-COMBINE-lab/ppi.bio. Many online academic servers are no longer available online and do not provide their source openly, resulting in the total loss of these servers (See Supplementary File S2 for a review of 22 such resources, 50% of which are no longer alive). By making the code public under an open-source license, we hope to enable others to run their own instance of the server in the event that https://ppi.bio is no longer online.

The process for making proteome-wide inferences on user-submitted amino acid sequence is as follows. Users input amino acid sequences which are then submitted to the PPI.bio server (Figure 5). The PPI.bio server in turn encodes the amino acid sequence using the user’s choice of either RAPPPID or INTREPPPID’s AWD-LSTM encoder to obtain a protein embedding. The embedding of the user-submitted protein is then paired with pre-computed embeddings of all proteins belonging to the proteome of the user’s choosing (currently, proteins for six organisms have been computed).

**Figure 5:**
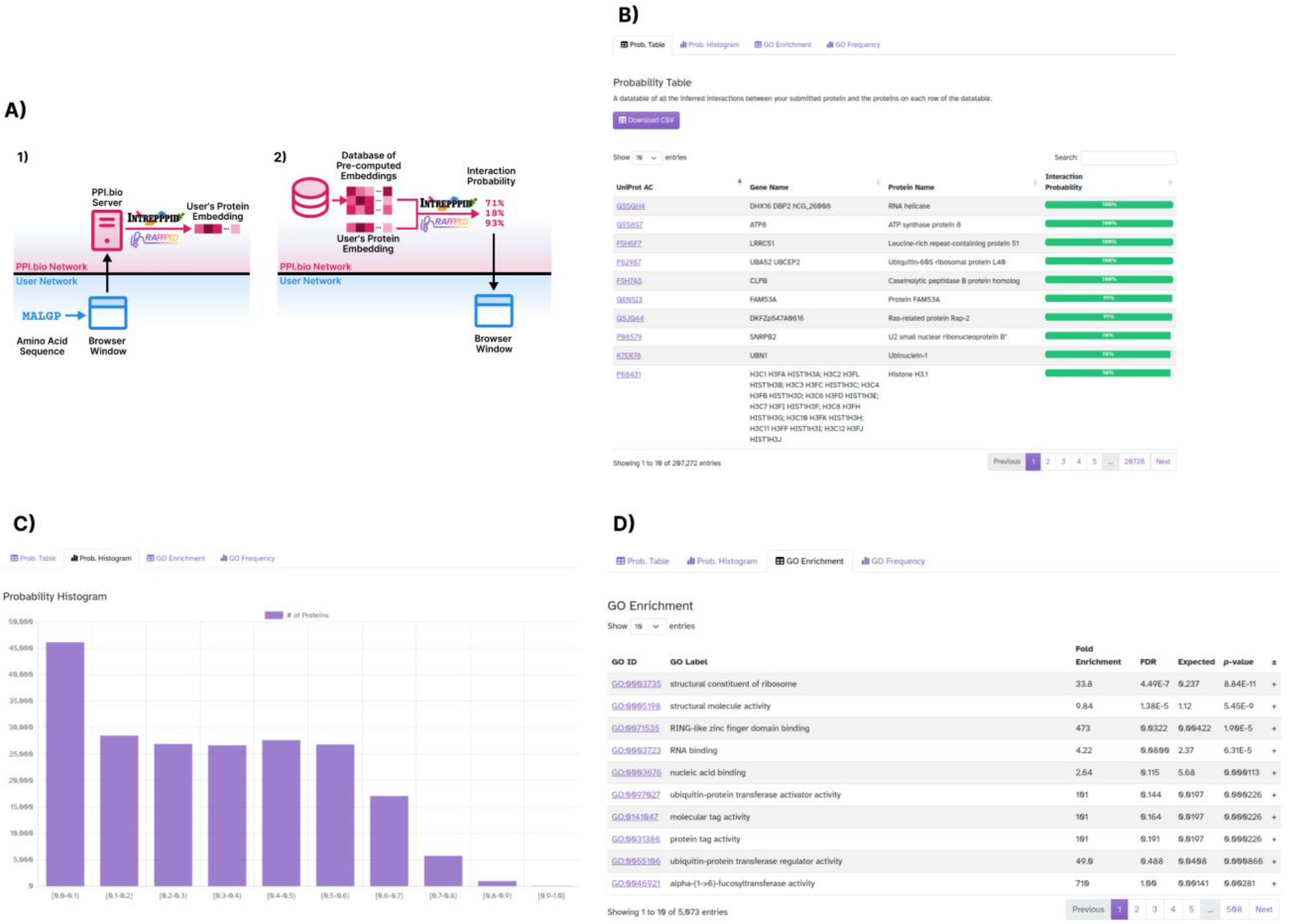
The webserver of PPI.bio supporting RAPPPID and INTREPPPID at whole proteome scale. **(A)** Overview of the operation of the PPI.bio PPI inference server. In the first stage (leftmost panel), amino acid sequences are encoded into protein embeddings using either RAPPPID or INTREPPPID, depending on the user’s preference. In the second stage (rightmost panel), pre-computed protein embeddings of a specified species proteome are loaded from a database. The classification head of either RAPPPID or INTREPPPID is then used to compute the probability of interaction of the user-specified protein and the pre-computed embeddings. Screenshot of the **(B)** data table of interaction probabilities, **(C)** probability histograms, and **(D)** GO enrichment analysis powered by PANTHERDB, as seen on PPI.bio after inferring proteome-wide interactions.

The probability of these pairs of proteins interacting are then inferred using RAPPPID or INTREPPPID’s classification head (based on the user’s choice), and returned to the user’s browser window.

PPI.bio also enables users to conduct GO enrichment analysis on the set of proteins that have an interaction probability greater than some user-specified threshold. PPI.bio uses the PANTHER online API (38) to calculate GO enrichment statistics. To illustrate the utility of this functionality, we chose at random a protein from the testing: the Mediator of RNA polymerase II transcription subunit 11, a product of the MED11 gene (UniProtKB Accession Q9P086). After computing the *H. sapiens* proteome-wide interactions of its reference sequence from UniProt, we conducted a GO enrichment analysis. Specifically, we analysed the enrichment among proteins which INTREPPPID inferred would interact with MED11 with a probability greater than 90%. We used the GO Molecular Function annotation dataset. We found that PPI.bio’s GO Enrichment analysis identified among its top enriched GO terms “RING-like zinc finger domain binding” (GO:0071535) to be 473 times enriched with a *p*-value of 1.90E-5 and “RNA binding” (GO:0003723) to be 4 times enriched with a *p*-value of 6.31E-5 (Figure 5C). This analysis correctly identifies that MED11 primarily interacts with proteins that bind RNA. Other notable enriched GO terms among inferred interactors include “structural molecule activity” (GO:0005198) at 9.84 fold enrichment with a *p*-value of 5.45E-9, and “protein-containing complex binding” (GO:0044877) at 2.42 fold enrichment with a *p*-value of 0.0337. This indicates that the analysis also correctly identifies the MED11 as being part of a larger complex.

## DISCUSSION

This study introduces three tools for researchers in their pursuit of better understanding PPIs: a cross-species method for PPI inference (INTREPPPID), a software package for creating strict PPI datasets (PPI Origami), and a web server for inferring PPIs using INTREPPPID and RAPPPID (PPI.bio). These tools and their code are all freely available under open licenses with complete documentation.

Through the adoption of the novel orthologous locality loss, INTREPPPID shows consistent gains over leading PPI inference methods in both the intra-species and cross-species task on strict PPI datasets. While INTREPPPID has a quintuplet network, there is no appreciable increase in trainable parameters compared to RAPPPID’s twin network construction due to the parameter sharing between encoders (2.20E5 and 1.88E5 parameters, respectively). By sharing parameters, INTREPPPID reduces both its GPU memory footprint as well as the risk of over-fitting via over-parameterization.

The orthologous locality task can be viewed through the lens of contrastive self-supervised learning (SSL) techniques like “SimCLR” or Time-Contrastive Networks (39, 40). Specifically, orthologous sequences can be thought of as multiple views of the same protein augmented by an evolutionary process in a way that is analogous to how SimCLR compares pairs of images which have been augmented by *e*.*g*.: cropping or blurring. In both instances, the model is learning invariance between entities which are similar in their desired feature space, but are significantly different in the input space. By creating a feature space that is invariant to the effects of speciation on amino acid sequences, we position our model to capture more interologues. While SimCLR only uses pairs of contrasting examples, SSL methods like Time-Contrastive Networks have adopted a triplet loss to more efficiently learn embeddings that better fit downstream tasks.

The final major contribution herein is PPI.bio, a web server for inferring PPIs using RAPPPID and INTREPPPID. PPI.bio infers interactions entirely on the CPU, greatly reducing server costs. Both RAPPPID and INTREPPPID facilitate this by allowing known protein sequences to be pre-computed, only requiring a forward pass of the shallow fully-connected classification network when the latent representation for both proteins has already been calculated. This allows for significant speed-ups when doing proteome-wide inferences.

As time passes, online services published in academic articles tend to go offline or otherwise become inaccessible. After conducting a search for the terms “protein-protein interaction portal” and “protein-protein interaction website” on the online academic search engine SemanticScholar.org and randomly sampling 23 articles which introduce online tools, 50% of the URIs either did not resolve or no longer offered the service introduced (Supplemental File S2). To help ensure that PPI.bio remains accessible for years to come, we’ve made the code that runs PPI.bio available under a free and open license. To help facilitate the deployment of PPI.bio, the server is designed to be easily deployed to the Platform as a Service (PaaS) service Heroku as well as the self-hosted alternative Dokku.

## SOFTWARE & DATA AVAILABILITY

INTREPPPID and the PPI.bio server are freely available under the Affero General Public License version 3.0. PPI Origami is licensed under the GNU General Public License version 3.0. They can be downloaded from https://github.com/Emad-COMBINE-lab/intrepppid, https://github.com/Emad-COMBINE-lab/ppi.bio, and https://github.com/Emad-COMBINE-lab/ppi_origami, respectively.

All code and datasets have been deposited in the Zenodo repository under record numbers 10652231, 10652229, 10652233, and 10594149.

## SUPPLEMENTARY MATERIAL

File S1: Contains supplementary Tables S2, S3, and S5. Supplementary Table S1 and S4 are provided as a separate .xlsx file (its caption is provided in File S1).

File S2: Contains the results of an analysis of 23 randomly sampled articles advertising online PPI-related tools and the online availability of said tools as of January 2024.

## AUTHORS’ CONTRIBUTIONS

A.E. and J.S. conceived the study, designed the project and the algorithm, and wrote the article. A.E. supervised the study. J.S. implemented the software, and performed the statistical analyses of the results. All authors read and approved the final article.

## FUNDING

This work was supported by grants from Natural Sciences and Engineering Research Council of Canada (NSERC) [RGPIN-2019-04460] (AE) and Canada Foundation for Innovation (CFI) JELF [project 40781]. This research was enabled in part by support provided by Calcul Québec (www.calculquebec.ca) and the Digital Research Alliance of Canada (alliancecan.ca).

## Conflict of Interest

The authors have not conflict of interest to declare.

## Notes

https://github.com/Emad-COMBINE-lab/intrepppid

https://github.com/Emad-COMBINE-lab/ppi.bio

https://github.com/Emad-COMBINE-lab/ppi_origami

https://zenodo.org/doi/10.5281/zenodo.10652231

https://zenodo.org/doi/10.5281/zenodo.10652229

https://zenodo.org/doi/10.5281/zenodo.10652233

https://zenodo.org/doi/10.5281/zenodo.10594149

### Competing Interest Statement

The authors have declared no competing interest.

## REFERENCES

1. Rolland, T., Taşan, M., Charloteaux, B., Pevzner, S.J., Zhong, Q., Sahni, N., Yi, S., Lemmens, I., Fontanillo, C., Mosca, R., et al. (2014) A proteome-scale map of the human interactome network. Cell, 159, 1212–1226.

2. Luck, K., Kim, D.-K., Lambourne, L., Spirohn, K., Begg, B.E., Bian, W., Brignall, R., Cafarelli, T., Campos-Laborie, F.J., Charloteaux, B., et al. (2020) A reference map of the human binary protein interactome. Nature, 580, 402–408.

3. Schoch, C.L., Ciufo, S., Domrachev, M., Hotton, C.L., Kannan, S., Khovanskaya, R., Leipe, D., Mcveigh, R., O’Neill, K., Robbertse, B., et al. (2020) NCBI Taxonomy: a comprehensive update on curation, resources and tools. Database, 2020, baaa062.

4. Szklarczyk, D., Kirsch, R., Koutrouli, M., Nastou, K., Mehryary, F., Hachilif, R., Gable, A.L., Fang, T., Doncheva, N.T., Pyysalo, S., et al. (2023) The STRING database in 2023: Protein–protein association networks and functional enrichment analyses for any sequenced genome of interest. Nucleic Acids Research, 51, D638–D646.

5. International Commission on Zoological Nomenclature (1999) International code of zoological nomenclature 4th ed. Ride, W.D.L., Cogger, H.G., Dupuis, C., Kraus, O., Minelli, A., Thompson, F.C., Tubbs, P.K. (eds) International Trust for Zoological Nomenclature, c/o Natural History Museum, London.

6. Park, Y. and Marcotte, E.M. (2012) Flaws in evaluation schemes for pair-input computational predictions. Nature Methods, 9, 1134–1136.

7. Dunham, B. and Ganapathiraju, M.K. (2021) Benchmark evaluation of protein–protein interaction prediction algorithms. Molecules, 27, 41.

8. Bernett, J., Blumenthal, D.B. and List, M. (2023) Cracking the black box of deep sequence-based protein-protein interaction prediction. bioRxiv.

9. Hamp, T. and Rost, B. (2015) More challenges for machine-learning protein interactions. Bioinformatics, 31, 1521–1525.

10. Marcotte, E.M., Pellegrini, M., Ng, H.-L., Rice, D.W., Yeates, T.O. and Eisenberg, D. (1999) Detecting protein function and protein-protein interactions from genome sequences. Science, 285, 751–753.

11. Walhout, A.J.M., Sordella, R., Lu, X., Hartley, J.L., Temple, G.F., Brasch, M.A., Thierry-Mieg, N. and Vidal, M. (2000) Protein interaction mapping in c. Elegans using proteins involved in vulval development. Science, 287, 116–122.

12. Matthews, L.R., Vaglio, P., Reboul, J., Ge, H., Davis, B.P., Garrels, J., Vincent, S. and Vidal, M. (2001) Identification of potential interaction networks using sequence-based searches for conserved protein-protein interactions or ‘interologs’. Genome Research, 11, 2120–2126.

13. Sharan, R., Suthram, S., Kelley, R.M., Kuhn, T., McCuine, S., Uetz, P., Sittler, T., Karp, R.M. and Ideker, T. (2005) Conserved patterns of protein interaction in multiple species. Proceedings of the National Academy of Sciences, 102, 1974–1979.

14. Cortes, C. and Vapnik, V. (1995) Support-vector networks. Machine Learning, 20, 273– 297.

15. Ben-Hur, A. and Noble, W.S. (2005) Kernel methods for predicting protein-protein interactions. Bioinformatics, 21, i38–i46.

16. Richoux, F., Servantie, C., Borès, C. and Téletchéa, S. (2019) Comparing two deep learning sequence-based models for protein-protein interaction prediction. arXiv:1901.06268 [cs, q-bio, stat].

17. Bromley, J., Bentz, J.W., Bottou, L., Guyon, I., Lecun, Y., Moore, C., Säckinger, E. and Shah, R. (1993) Signature verification using a ‘siamese’ time delay neural network. International Journal of Pattern Recognition and Artificial Intelligence, 07, 669– 688.

18. Baldi, P. and Chauvin, Y. (1993) Neural networks for fingerprint recognition. Neural Computation, 5, 402–418.

19. Koch, G.R., Zemel, R. and Salakhutdinov, R. (2015) Siamese neural networks for one-shot image recognition. In Proceedings of the 32^nd^ international conference on machine learning.Vol. 37.

20. Chen, M., Ju, C.J.-T., Zhou, G., Chen, X., Zhang, T., Chang, K.-W., Zaniolo, C. and Wang, W. (2019) Multifaceted protein–protein interaction prediction based on siamese residual RCNN. Bioinformatics, 35, i305–i314.

21. Cho, K., Merrienboer, B. van, Gulcehre, C., Bahdanau, D., Bougares, F., Schwenk, H. and Bengio, Y. (2014) Learning phrase representations using RNN encoder-decoder for statistical machine translation. 10.48550/arXiv.1406.1078.

22. Szymborski, J. and Emad, A. (2022) RAPPPID: Towards generalizable protein interaction prediction with AWD-LSTM twin networks. Bioinformatics, 38, 3958– 3967.

23. Merity, S., Keskar, N.S. and Socher, R. (2017) Regularizing and optimizing LSTM language models. arXiv:1708.02182 [cs].

24. Kudo, T. and Richardson, J. (2018) SentencePiece: A simple and language independent subword tokenizer and detokenizer for neural text processing. In Blanco, E., Lu, W. (eds), Proceedings of the 2018 conference on empirical methods in natural language processing: System demonstrations. Association for Computational Linguistics, Brussels, Belgium, pp. 66–71.

25. Kudo, T. (2018) Subword regularization: Improving neural network translation models with multiple subword candidates. arXiv.org.

26. Misra, D. (2020) Mish: A self regularized non-monotonic activation function. arXiv:1908.08681 [cs, stat].

27. Zhang, X., Yu, F.X., Kumar, S. and Chang, S.-F. (2017) Learning spread-out local feature descriptors. In 2017 IEEE international conference on computer vision (ICCV). IEEE, Venice, pp. 4605–4613.

28. Altenhoff, A.M., Train, C.-M., Gilbert, K.J., Mediratta, I., Mendes de Farias, T., Moi, D., Nevers, Y., Radoykova, H.-S., Rossier, V., Warwick Vesztrocy, A., et al. (2021) OMA orthology in 2021: Website overhaul, conserved isoforms, ancestral gene order and more. Nucleic Acids Research, 49, D373–D379.

29. Schroff, F., Kalenichenko, D. and Philbin, J. (2015) FaceNet: A unified embedding for face recognition and clustering. In 2015 IEEE conference on computer vision and pattern recognition (CVPR).pp. 815–823.

30. Forslund, K. (2011) The relationship between orthology, protein domain architecture and protein function.

31. Fang, G., Bhardwaj, N., Robilotto, R. and Gerstein, M.B. (2010) Getting started in gene orthology and functional analysis. PLoS Computational Biology, 6, e1000703.

32. Suzek, B.E., Wang, Y., Huang, H., McGarvey, P.B., Wu, C.H. and UniProt Consortium,the (2015) UniRef clusters: A comprehensive and scalable alternative for improving sequence similarity searches. Bioinformatics, 31, 926–932.

33. Consortium, T.U., Bateman, A., Martin, M.-J., Orchard, S., Magrane, M., Ahmad, S., Alpi, E., Bowler-Barnett, E.H., Britto, R., Bye-A-Jee, H., et al. (2023) UniProt: The universal protein knowledgebase in 2023. Nucleic Acids Research, 51, D523–D531.

34. Ben-Hur, A. and Noble, W.S. (2006) Choosing negative examples for the prediction of protein-protein interactions. BMC Bioinformatics, 7, S2–S2.

35. Sledzieski, S., Singh, R., Cowen, L. and Berger, B. (2021) D-SCRIPT translates genome to phenome with sequence-based, structure-aware, genome-scale predictions of protein-protein interactions. Cell Systems, 12, 969–982.e6.

36. Li, Y. and Ilie, L. (2017) SPRINT: Ultrafast protein-protein interaction prediction of the entire human interactome. BMC Bioinformatics, 10.1186/s12859-017-1871-x.

37. McInnes, L., Healy, J. and Melville, J. (2020) UMAP: Uniform manifold approximation and projection for dimension reduction. arXiv:1802.03426[stat.ML].

38. Thomas, P.D., Ebert, D., Muruganujan, A., Mushayahama, T., Albou, L. and Mi, H. (2022) PANTHER : Making genome-scale phylogenetics accessible to all. Protein Science, 31, 8–22.

39. Chen, T., Kornblith, S., Norouzi, M. and Hinton, G. (2020) A simple framework for contrastive learning of visual representations.

40. Sermanet, P., Lynch, C., Chebotar, Y., Hsu, J., Jang, E., Schaal, S. and Levine, S. (2018) Time-contrastive networks: Self-supervised learning from video.

41. Alanis-Lobato, G., Möllmann, J.S., Schaefer, M.H. and Andrade-Navarro, M.A. (2020) MIPPIE: The mouse integrated protein–protein interaction reference. Database: The Journal of Biological Databases and Curation, 2020, baaa035.

42. Kusari, M., Dey, L. and Mukhopadhyay, A. (2022) ChikvInt: A chikungunya virus–host protein–protein interaction database. Letters in Applied Microbiology, 74, 992– 1000.

43. Kalman, Z.E., Dudola, D., Mészáros, B., Gáspári, Z. and Dobson, L. (2022) PSINDB: The postsynaptic protein–protein interaction database. Database, 2022, baac007.

44. Wright, L. and Demeure, N. (2021) Ranger21: a synergistic deep learning optimizer.

45. Loshchilov, I. and Hutter, F. (2019) Decoupled Weight Decay Regularization. arXiv:1711.05101 [cs, math].

46. Iyer, N., Thejas, V., Kwatra, N., Ramjee, R. and Sivathanu, M. (2021) Wide-minima Density Hypothesis and the Explore-Exploit Learning Rate Schedule. 10.48550/arXiv.2003.03977.

