## Supplemental File S1 for "INTREPPPID - An Orthologue-Informed Quintuplet Network for Cross-Species Prediction of Protein-Protein Interaction"

### Supplementary Materials

#### Tables

**Table S1** is provided as a separate .xlsx spreadsheet file.

Its caption is as follows:

> Statistics of each dataset used in this study. Datasets are listed by source organism and random seed used to generate the random splits.

**Table S2** - Species for which PPI data from STRING was used in the analyses herein.

Patristic distance is a measure of the divergence of two lineages. Patristic distances were computed using DendroPy and data from iTOL (1,2).

| Scientific Name | Common Name | NCBI Taxon ID | Patristic Distance from *H. sapiens* |
|---|---|---|---|
| *Homo sapiens* | Human | 9606 | 0.000 |
| *Mus musculus* | House mouse | 10090 | 0.0455 |
| *Danio rerio* | Zebrafish | 7955 | 0.156 |
| *Drosophila melanogaster* | Fruit fly | 7227 | 0.358 |
| *Caenorhabditis elegans* | Nematode | 6239 | 0.455 |
| *Arabidopsis thaliana* | Thale cress | 3702 | 0.693 |
| *Saccharomyces cerevisiae* | Baker’s yeast | 4932 | 0.734 |

**Table S3** - Hyperparameters used to train INTREPPPID in all experiments.

| Hyperparameter | Value |
|---|---|
| Batch Size | 80 |
| Embedding Size | 64 |
| Number of RNN Layers | 2 |
| RNN Drop-connect Rate | 0.3 |
| Embedding Drop-out Rate | 0.3 |

**Table S4** is provided as a separate .xlsx spreadsheet file.

Its caption is as follow:

> Metrics of various PPI inference methods trained on *H. sapiens* data, and tested on various species. Each model was tested on a total of twenty-four dataset: four different random testing splits for each six different organism. The metrics reported here are averaged over all four datasets, with the standard deviation between the four datasets also reported. The first worksheet (AUROC) reports the area-under-the-receiver-operator-curve (AUROC). The second worksheet (AP) reports the average-precision. The third worksheet (F150) reports the F1 measure with a 50% threshold. The fourth worksheet (MCC50) reports the Matthew's correlation coefficient (MCC) with a 50% threshold. The fifth and final worksheet (Brier) reports the Brier score. Higher values indicate better performance for all metrics but the Brier score, where the inverse is true.

**Table S5** - Performance of INTREPPPID trained on *H. sapiens* datasets with scrambled and not scrambled orthologue information. Bold-face indicates the best value for each metric.

| Scramble Status | Test AUROC | Test AP | Test MCC@50% | Test Brier Score |
|---|---|---|---|---|
| Not Scrambled | **0.761** | **0.737** | **0.378** | **0.198** |
| Scrambled | 0.524 | 0.527 | 0.017 | 0.314 |

#### References


1. Letunic,I. and Bork,P. (2021) Interactive Tree Of Life (iTOL) v5: an online tool for phylogenetic tree display and annotation. *Nucleic Acids Research*, 49, W293–W296.

2. Sukumaran,J. and Holder,M.T. (2010) DendroPy: a Python library for phylogenetic computing. *Bioinformatics*, 26, 1569–1571.
